# Evaluating Preservation Techniques for Long-Term Stability of 3D Bioprinted Liver Scaffolds

**DOI:** 10.64898/2026.03.11.711081

**Authors:** Mrunmayi Gadre, Kirthanashri S Vasanthan

## Abstract

Three-dimensional (3D) bioprinted liver scaffolds offer a promising platform for drug screening, disease modelling, and regenerative medicine, yet their broader adoption is limited by the absence of robust post-fabrication preservation strategies. This study aimed to evaluate the impact of −80°C (deep freezer) preservation and evaluate the structural integrity and hepatic functionality of GelMA–decellularized liver extra cellular matrix (dECM)–based 3D bioprinted liver scaffolds. Bioinks were formulated using synthesized GelMA and solubilized rat liver dECM, and 3D scaffolds were fabricated via extrusion bioprinting into rectilinear grid scaffolds. The 3D scaffold preservations was performed by immersion into two different medium (the culture DMEM media and the other FBS-DMSO cocktail) was evaluated using MTT viability assay, and albumin assay. Preserved 3D bioprinted scaffolds retained overall architecture and cell distribution in the FBS-DMSO cocktail demonstrated by the live dead assay. Together, the data demonstrate that −80°C storage can maintain the basic cell viability (∼80%) and a substantial fraction of liver-specific functionality in 3D bioprinted scaffolds but also highlight sensitivity to preservation-induced injury. These findings underscore the need for further optimization of cryoprotectant formulations and freezing protocols tailored to 3D bioprinted liver scaffolds, and provide a foundational framework for developing ready-to-use, cryopreserved 3D liver models for translational applications.

## 1. INTRODUCTION

Over the last few decades there has been rapid evolution in the field of tissue engineering and one such technique is 3D bioprinting. The 3D bioprinting is precise deposition of cell-laden bioinks in layer-by-layer manner. The constructs fabricated by 3D bioprinting provide a tailored architecture and the application have a wide application like *in vitro* modeling, drug testing platforms and regenerative therapies that can be developed patient specific [Sharma R et al. 2023; Mally D et al. 2025]. The 3D bioprinted liver scaffolds have been arisen to be an important platform with wide applications like drug screening platform, disease modelling and regenerative medicine. **Figure 1** streamlines the overall methodology applied to construct the 3D *in vitro* liver scaffold. These 3D *in vitro* models are more relevant physiologically over the traditional 2D cultures and simple spheroids [Xie R et al. 2024; Zhao Y et al. 2025; Ma Y et al. 2025]. However, the benchtop procedures are robust and time consuming for production and maintaining the replicating results in each batch becomes very challenging [Kim MK et al. 2025]. The quality of products to be categorised as clinical grade depends on the ability of the product’s storage, transportation and no significant loss in the functionality and structural integrity. The preservation of 3D *in vitro* models in long-term has become a central bottleneck in tissue engineering. The lack of availability of reliable preservation procedures, a researcher has to 3D bioprint *in vitro* models for immediate utilization. Limited scalability, batch to batch variability and lack of preservation techniques, leads to the loss in the integration of such products into industrial pipelines [Freitas-Ribeiro S et al. 2025; Budharaju H et al. 2024; Agarwal T et al. 2025].

**Figure 1:**
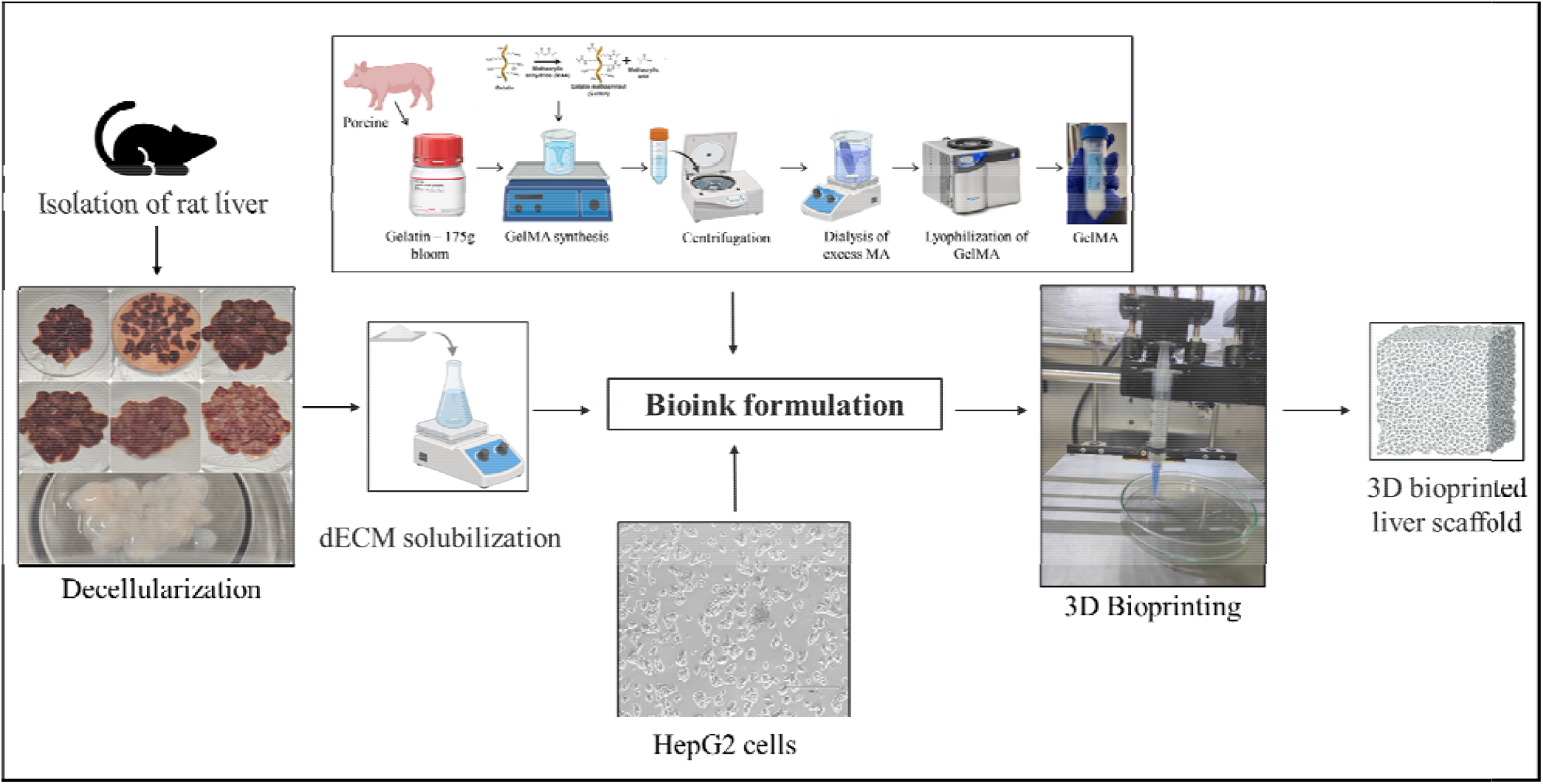
The illustration depicts the overview of the experimental workflow for fabricating decellularized liver dECM–GelMA composite bioprinted scaffolds by extrusion-based 3D bioprinter

Since the development in technology in the field of tissue engineering, the entire focus has been in the optimization of the biomaterials, fabricating techniques and sourcing of cells. There was less or no emphasis on the post-fabrication (storage and preservation practice) of 3D *in vitro* models [Budharaju H et al. 2024; Ziani K et al. 2025]. The available conventional methods of *in vitro* preservation involves various techniques are summarised in **table 1**. As per the best of authors knowledge, there are no reported method in literature that has been employed in the preservation of the 3D bioprinted scaffolds and hence maintaining the cell viability in 3D bioprinted scaffold is an unique challenges for cell preservation. The 3D scaffolds typically contains biomaterials, various crosslinkers, biological components (dECM/cytokines/growth factors) and cells, which makes the preservation of 3D scaffolds highly sensitive. There can be factors that damage the 3D scaffolds as they are highly hydrated and post preservation (freezing/thawing cycles), leads to structural damage resulting in loss of mechanical strength [Sever M et al. 2024]. Ideally 3D scaffolds should not only mimic physiological microenvironment but also maintain the functionality and biological features [Ziani K et al. 2025]. Recent advancements in tissue engineering suggests that preservation of 3D *in vitro* models might be feasible but require a systematic evaluation. The field also currently lacks a standardized and evidence-based guidelines on the best method to preserve the 3D bioprinted scaffolds [Budharaju H et al. 2024, Moon et al. 2025].

**Table 1:**
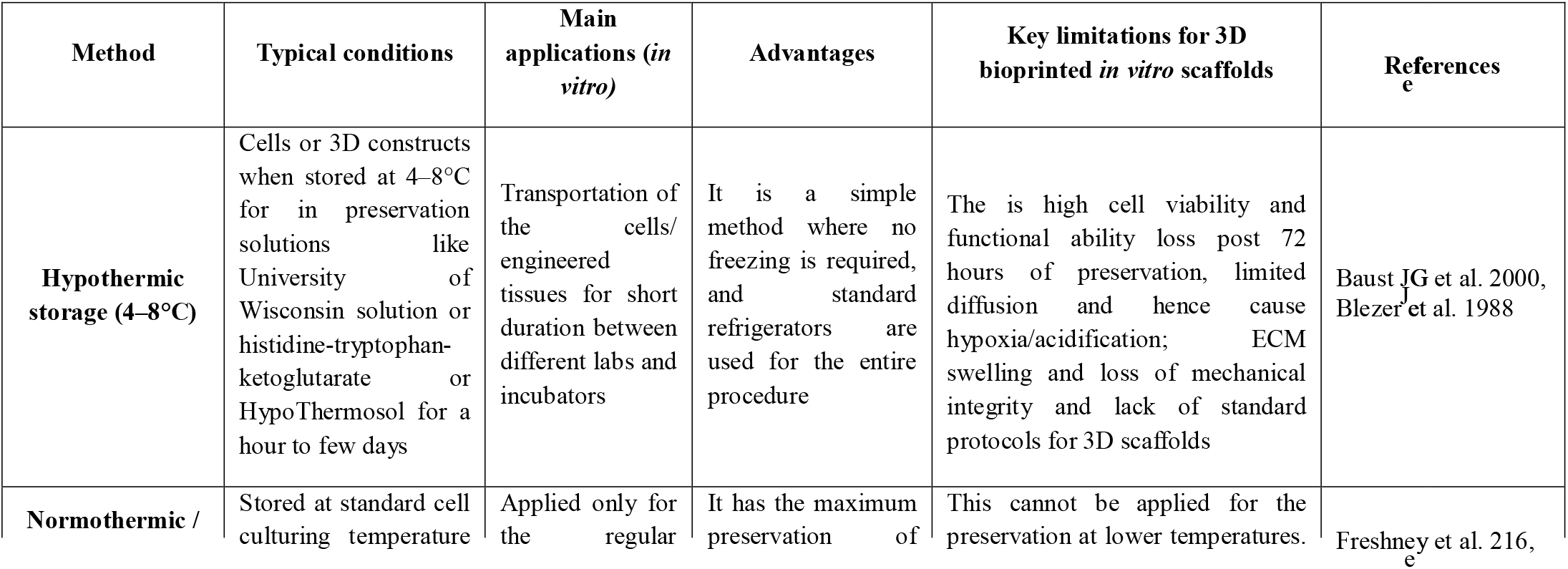
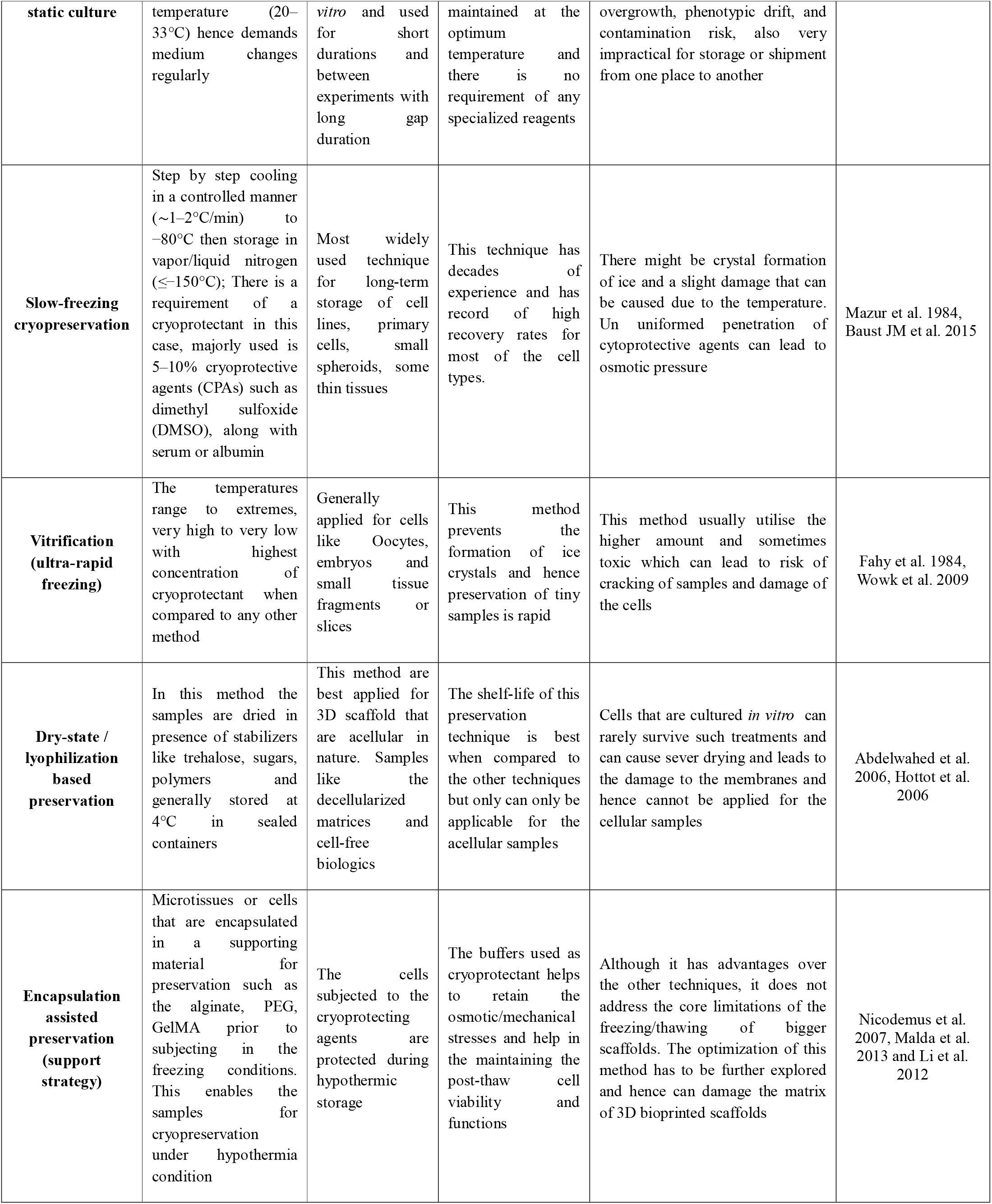
Different types of conventional *in vitro* preservation methods and drawbacks in preservation of 3D bioprinted scaffolds.

In this study, we have demonstrated that the preservation techniques for 3D bioprinted liver scaffolds are systematically evaluated and maintains liver-specific functionality post preservation at -80°C for defined time period (day 15, 30, 45, 60 and 90). **Figure 2** illustrates three different stages involved in 3D bioprinting process, including; pre-bioprinting stage (Computer aided designing is involved to generate G-codes for the 3D models); bioprinting stage (bioink is formulated and 3D bioprinted); and post-bioprinting stage (3D bioprinted scaffolds are processed for *in vitro*/*in vivo* study).

**Figure 2:**
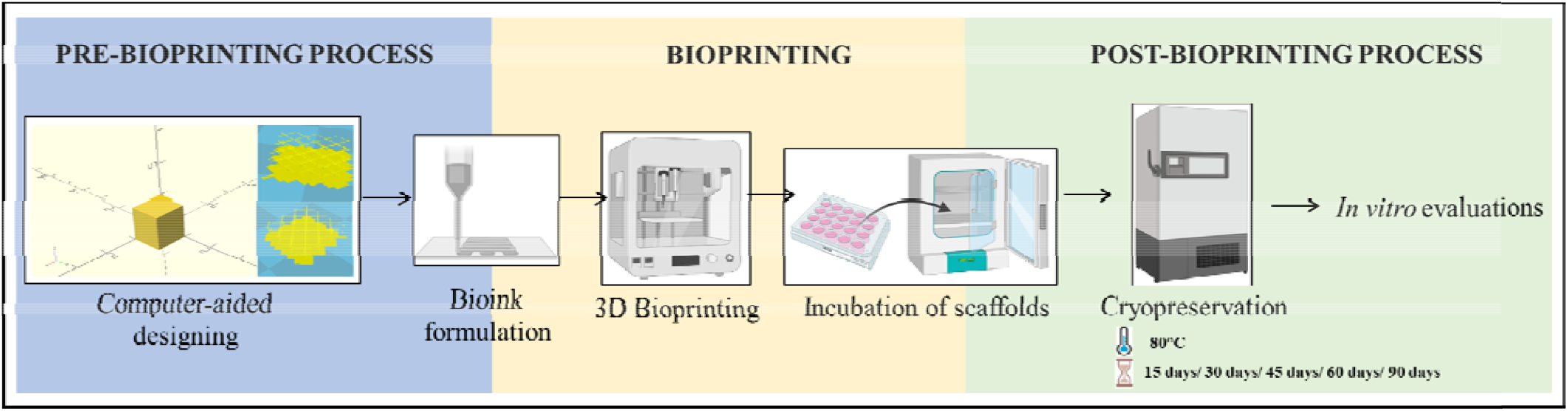
The illustration depicts the different stages involved in 3D bioprinting process

Building on the recent advances in liver tissue engineering and organ preservation, long-term stability of 3D bioprinted scaffolds was assessed for cell viability and expression of hepatic functional markers. The prime objective of the study is to identify preservation strategies that can support ready-to-use (from freezer to culture), 3D bioprinted liver scaffolds which can be applicable for drug testing and disease modelling. Developing such 3D bioprinted scaffold will ease the researchers and preserve the platforms for validation of drugs for future use.

Ultimately, by establishing robust preservation protocols and standardising the protocols can improve the reproducibility, scalability, and translational potential of 3D *in vitro* models for research use.

With the advancement in tissue engineering, decellularized extra cellular matrix (dECM) has gained promising results to mimick organs *in vitro* [Kasturi et al. 2024; Gadre M et al. 2024]. Scientists are utilizing dECM in bioink for bioprinting to generate *in vivo* like 3D bioprinted scaffolds for varied application. The scaffolds produced using 3D bioprinting technique has been established by the group in previous study [Vidhi et al. 2023; Gadre M et al. 2025]. The goal of this study is cryopreserve 3D bioprinted scaffolds by integrating Gelatin methacylamide (GelMA) combined with rat liver-derived dECM, which helps provide the essential native signals. Besides these two ingredients, the bioink also comprised of crosslinking agents and HepG2 cells creating environment that helps in hepatic cell function.

## 2.0 MATERIALS AND METHODOLOGY

### 2.1 Materials

The cell culture experiments were sourced from Himedia, India conducted using Dulbecco’s Phosphate-Buffered Saline (DPBS) and media Dulbecco’s Modified Eagle’s Medium (DMEM). Fetal bovine serum (FBS) was supplied by Gibco. Essential reagents, including trypsin, antibiotics (penicillin/streptomycin), methacrylic anhydride, formaldehyde and trizol were purchased from Sigma Aldrich. The Abcam supplied with the Albumin (BCG) Assay Kit [ab235628]. Chemicals such as isopropanol, papain, chloroform, sodium dodecyl sulphate (SDS), isoamyl alcohol and phenol were purchased from Sisco Research Laboratories, India. Other reagents like formaldehyde and ethylenediamine tetraacetic acid (EDTA) came from Qualigens, India. Phosphate-Buffered Saline (PBS), Gelatin type A (Bloom strength 175 g, porcine source), glutaraldehyde from Sigma Aldrich, India. Live dead viability assay kit [L3224] was acquired from Thermo Fisher Scientific, India. Other essential chemicals such as 3- (4,5-dimethylthiazol-2-yl)-2,5-diphenyltetrazolium bromide (MTT) and Dimethyl sulfoxide (DMSO) along with consumables like T25 and T75 cell culture flasks were brought from Himedia, India. Remaining consumables like the 6-well plates, 24 well-plates and 96-well plates, 15- and 50-mL centrifuge tubes were procured from Abdos life sciences, India.

### 2.2 Ethical approval for animal studies

All the protocols related to the animals was performed with approval of Institutional Animal Ethics Committee (IAEC), protocol number: IAEC/KMC/121/2022, at Manipal Academy of Higher Education, Manipal and under strict adherence to the ARRIVE guidelines, all the protocols including animals were carried out to ensure ethical and humane treatment throughout the research process.

### 2.3 METHODOLOGY

#### 2.3.1. The Cell Culture technique

The Human hepatocellular carcinoma cell culture (HepG2) which was utilised for the entire study was acquired from National Centre for Cell Science, India. Cells under complete culture conditions were cultivated in DMEM which was supplemented with 10% (v/v) fetal bovine serum (FBS) along with 1% (v/v) penicillin/streptomycin. The cells were sustained in a humidified incubator at 37°C with 5% CO□ to ensure optimal growth conditions.

#### 2.3.2. Computer-Aided Design (CAD) for 3D Bioprinting

The 3D models were generated using the CAD designed according to the study, the application openscad was used for generating G-code and STL file. All the parameters for the 3D model such as the dimensions, intended layers and the speed of the bioprinter and the pressure was specified.

#### 2.3.3. Bioink preparation

The GelMA was synthesized in house by using one pot method protocol [Vidhi et al. 2023]. Briefly, gelatin was dissolved in PBS (pH 7.4), along with crosslinker MA under constant stirring. The un-crosslinked MA was dialyzed against double distilled water by dialysis membrane and content was then lyophilized. GelMA along with rat liver dECM was utilized to formulate the bioink. Male Wistar rats, were carefully anesthetized and livers were carefully excised. Using chemical method (EDTA and SDS), decellularization was performed (60 mL per concentration was injected at 8–10 sites) and the obtained dECM was characterized for gold standard. HepG2 cells were cultured under sterile conditions and approximately 50,000 cells were used in the bioink formulation per scaffold, the resulting cell-laden bioink was filled in sterile syringe and 3D bioprinted [Gadre M et al. 2025].

#### 2.3.4. Cryopreservation of 3D bioprinted scaffolds

The general method for cryopreservation used for *in vitro* 2D culture is storages of cell pellet in liquid nitrogen with the cocktail of FBS and DMSO. The protocol applied in this study is storage of 3D bioprinted scaffolds in -80°C. **Figure 3** depicts the overall methodology of cryopreservation of the 3D bioprinted scaffolds. Post 3D bioprinting, 3D bioprinted scaffolds were incubated for 7 days at 37°C in CO_2_ incubator to allow the cell proliferation. The 3D bioprinted scaffolds were subjected to cryoprotectant; one group in which the 3D scaffolds had emerged into the culture DMEM media and the other had emerged into the FBS-DMSO cocktail in the ratio of 9:1. The cryovials were then stored for period of 15 days/ 30 days/ 45 days/ 60 days and 90 days for long-term preservation study. On each respective time point mentioned, the cryovials were removed and thawed to segregate the 3D bioprinted scaffolds from cryoprotectant. The 3D bioprinted scaffolds were then stored for a period of 48 hours at 37°C, then they were subjected to the *in vitro* evaluations.

**Figure 3:**
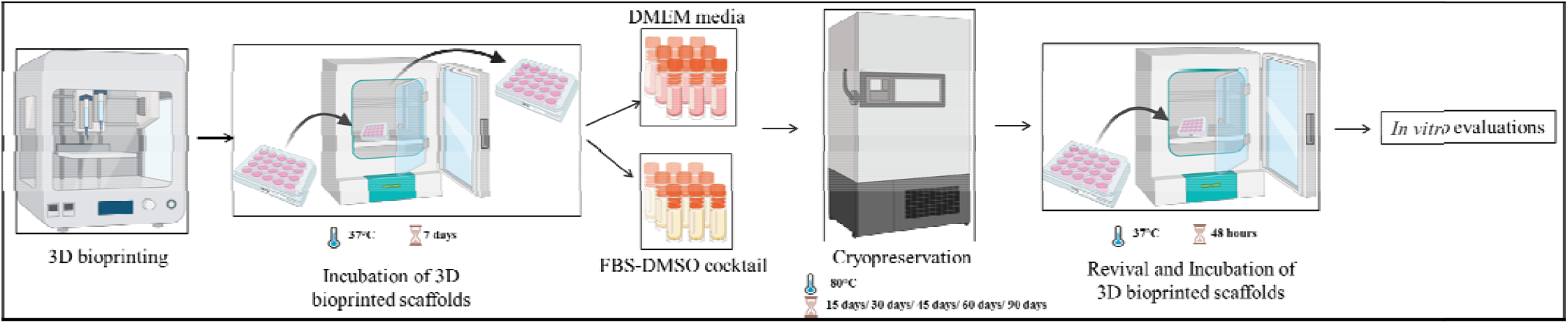
The image describes the methodology used for cryopreservation during this study

#### 2.3.5. Characterization of 3D bioprinted scaffolds post cryopreservation

##### 2.3.5.1. Metabolic Assay

Cell survival at each time point was evaluated by subjecting to 0.5% MTT solution and incubated for 4 hours **figure 4A**. The formazan crystals were dissolved in 300 μL of DMSO and at three different absorbance wavelength was measured (570 nm, 590 nm, and 630 nm) in microplate reader and was calculated cell metabolic activity with the reference of standard curve [Perez MG et al. 2017].

**Figure 4:**
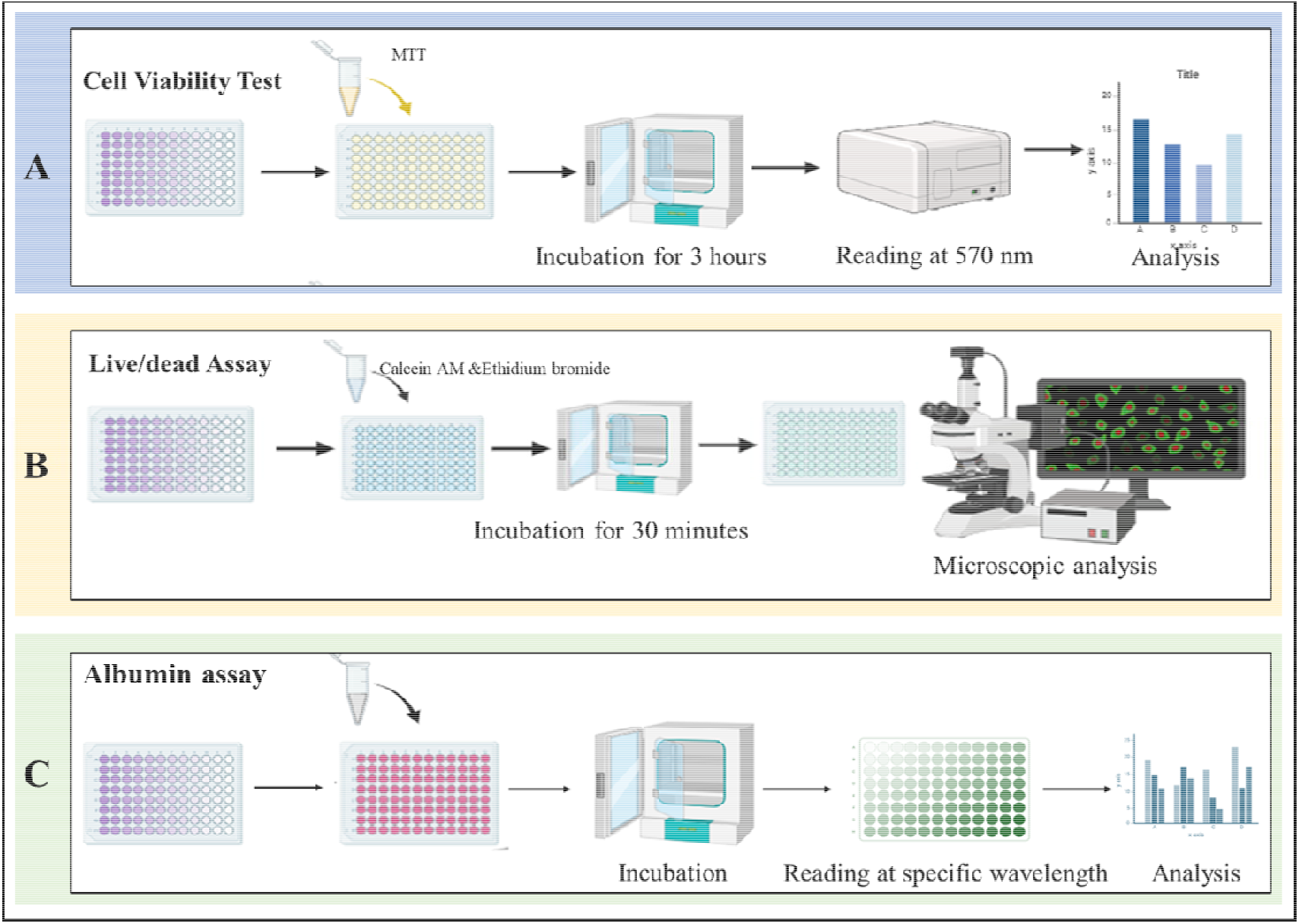
The graphical diagram represents the methodology employed in the study; (A) Cell viability assessed by MTT assay, (B) Cell viability assessed by live/dead assay and (C) Functional test (albumin assay) assessed by calorimetric method

##### 2.3.5.2. Live/Dead Assay

Cell survival by live dead assay at each time point was performed, strictly rendering to instructions provided by the manufacturer **figure 4B**. Briefly, a working solution was prepared by diluting 0.2 μM Calcein AM in 1 mL DPBS and 0.4 μM EthD-1 in the same solution [Dupont K et al. 2010]. Post incubation for 45 minutes in dark condition, the bioprinted scaffolds were observed using an EVOS M5000 fluorescence microscope (Invitrogen, India).

##### 2.3.5.3. Efflux Media Collection and Biochemical Measurements

The spent culture media were collected from 3D bioprinted scaffolds on 15 days/ 30 days/ 45 days/ 60 days and 90 days. Key liver secretion concentration (Albumin) were measured using commercially available kit-based methods as per manufacturer’s instructions **figure 4C**. The concentration was determined via a colorimetric kit assay utilizing Bromocresol Green (BCG), which forms a measurable chromophore with albumin at 620 nm. Briefly, 50 μL of sample was added to working solution (100 μL) and incubated for 25 minutes, and absorbance were recorded at 620 nm. Albumin concentrations were then calculated with the reference of standard curve [Poli V et al. 2023].

## 3.0 RESULTS AND DISCUSSION

### 3.1. Characterization of 3D bioprinted scaffolds post cryopreservation

The 3D bioprinted scaffolds that were standardized and characterized in the previous study was carried forward to this study using the same parameters and combinations. The 3D bioprinted scaffolds were proven to exhibit liver-specific functions hence were further applied to study the cryopreservation conditions.

### 3.2 Cell Viability Assay

Cell viability was evaluated on cryopreserved 3D bioprinted scaffolds at Day 0, 15, 30, 45, 60 and 90. The MTT was performed before the preservation (day 0) and post revival (day 15, 30, 45, 60 and 90). Post retrieval all the 3D bioprinted scaffolds were incubated for 48 hours to initiate the cell cycle [Huang G et al. 2024]. There was observed to be a consistent decline in viability of cells with increase in the incubation period (day 15, 30, 45, 60 and 90) on the 3D bioprinted scaffolds preserved in with DMEM media **(Figure 5A**). While 3D bioprinted scaffolds preserved in the FBS-DMSO cocktail demonstrated no decrease in cell viability over the period of time (day 15, 30, 45, 60 and 90) **(Figure 5B)**. This primarily indicates that the cocktail worked better than the DMEM media for the preservation of the 3D bioprinted scaffolds. DMSO in combination with FBS has resulted in reduced ice damage and osmotic shock leading to higher cell viability to maintain the functionality [Awan M et al. 2020].

**Figure 5:**
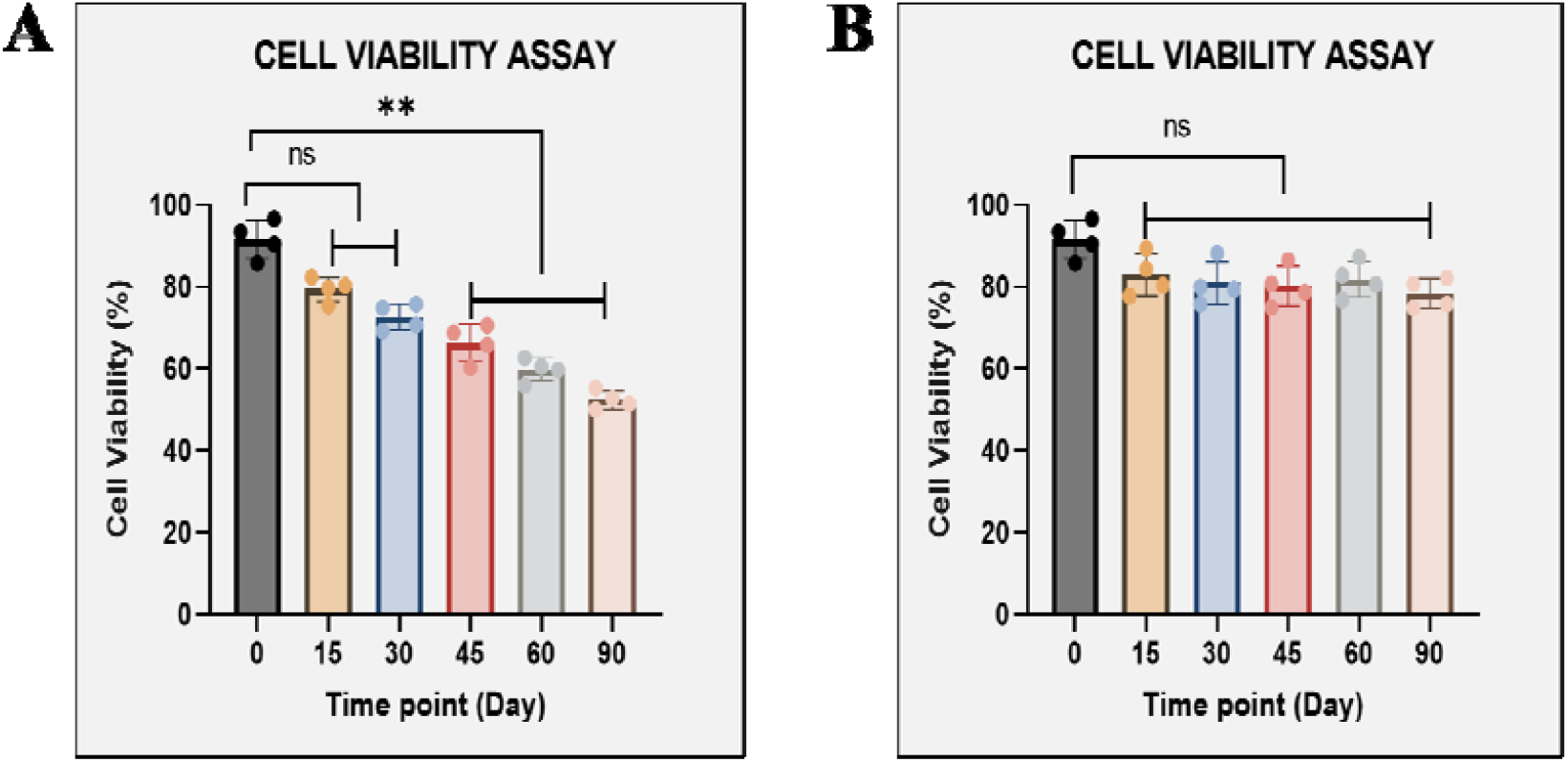
Cell viability analysis via MTT (A) 3D bioprinted scaffolds stored in DMEM media and (B) 3D bioprinted scaffolds stored in the DMSO and FBS cocktail ; Data presented as mean ± SD (n = 4) statistically significant ** : p<0.01 and ns: non-significant

Cryopreservation of cell-laden 3D scaffolds in DMEM at -80°C failed to maintain cell viability likely due to the absence of cryoprotective agents, resulting in uncontrolled ice crystal formation, osmotic stress, and mechanical disruption of both cellular membranes and scaffold microarchitecture. Additionally, non-uniform cooling rates, due to the absence of cryoprotectant lead to limited solute diffusion within the 3D matrix further exacerbated cell injury, while storage at -80°C may have promoted ice recrystallization and delayed apoptotic responses during post-thaw recovery [Ziani K et al. 2025].

As outlined in **figure 5B**, 3D GelMA–dECM scaffolds maintained approximately 80% of cell viability at all time point when preserved in the FBS-DMSO cocktail at (-80°C).

Temperature controlled cryoprintng is the reported technique for cryopreservation of 3D scaffolds, where the bioink deposition is performed on a descending, temperature-regulated freezing plate immersed within a cooling bath, which enables precise control of the nozzle temperature throughout the printing process [Warburton L at al. 2023]. This process has been reported to have cell viability of only 71±7.47%, whereas we claim that 3D bioprinted scaffolds upon preservation at -80°C with DMSO-FBS cocktail demonstrated higher cell viability of upto 81±2.59%.The efficacy of cell viability has been drastically improved and help the ready to use 3D bioprinted scaffold.

#### 3.3 Live/Dead Assay

Live/Dead (LD) staining was performed on each time point (day 15, 30, 45, 60 and 90) on 3D bioprinted scaffolds as shown in **Figure 6, which represents** a time-dependent decline in the live cells. Corresponding to the cell viability analysis using MTT **(Figure 5A&B)** similar trends were observed for LD assays that strengths our finding on the preservation technique and cell storage cocktail. **Figure 7 demonstrates that 3D bioprinted scaffold**-maintained cell viability throughout the time points by having consistent number of live cell fraction showing green fluorescence. This confirms that the standard cryoprotectant used for 2D cell culture also demonstrates cyro and cyto protectant behaviour on the 3D bioprinted scaffolds.

**Figure 6:**
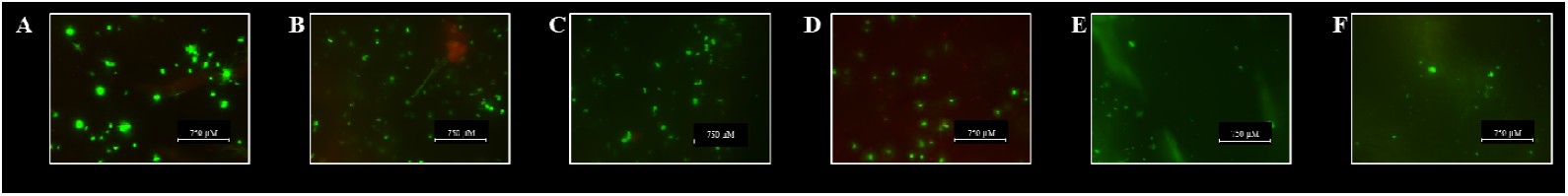
Representative fluorescence microscopy images of live (green) and dead (red) cells grown on 3D bioprinted scaffolds stored in DMEM media (A) Day 0 (B) Day 15 (C) Day 30 (D) Day 45 (E) Day 60 (F) Day 90 at 10X magnification

**Figure 7:**
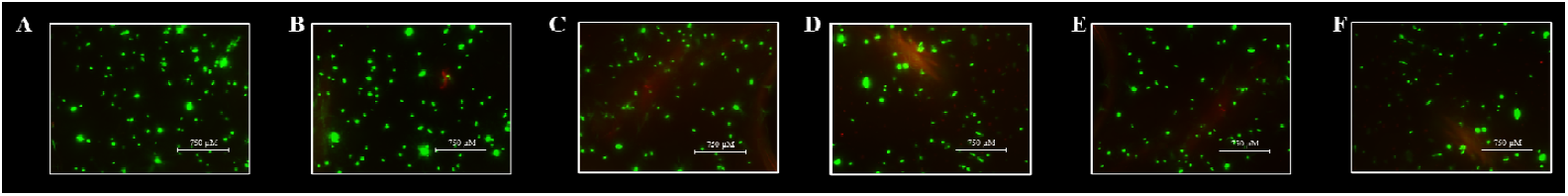
Representative fluorescence microscopy images of live (green) and dead (red) cells grown on 3D bioprinted scaffolds stored in DMSO and FBS cocktail (A) Day 0 (B) Day 15 (C) Day 30 (D) Day 45 (E) Day 60 (F) Day 90 at 10X magnification

### 3.4 Biochemical Measurement

Liver-specific function of the 3D bioprinted scaffolds was evaluated by assessing secretory (albumin) in the media spent **(Figure 8)**. The secretion of albumin in the DMEM medium showed significant decline at all the time point, indicating severe loss in the functionality of the 3D bioprinted scaffolds **(Figure 8A)**, while albumin secretion in DMSO-FBS cocktail showed constant range in all the time points **(Figure 8B)**. Albumin secretion in other 3D bioprinted scaffolds on day 7 were found to have secretion of ∼4-6 μg/mL Although the hepatic functionality in the 3D GelMA-dECM scaffolds were only retained in the FBS-DMSO cocktail, this leaves a strengthening and translational potential for researching more methods of preserving the 3D *in vitro* scaffolds. Although further assays and gene expression have to be carried out, this primary data supports the preservation of the 3D bioprinted scaffolds.

**Figure 8:**
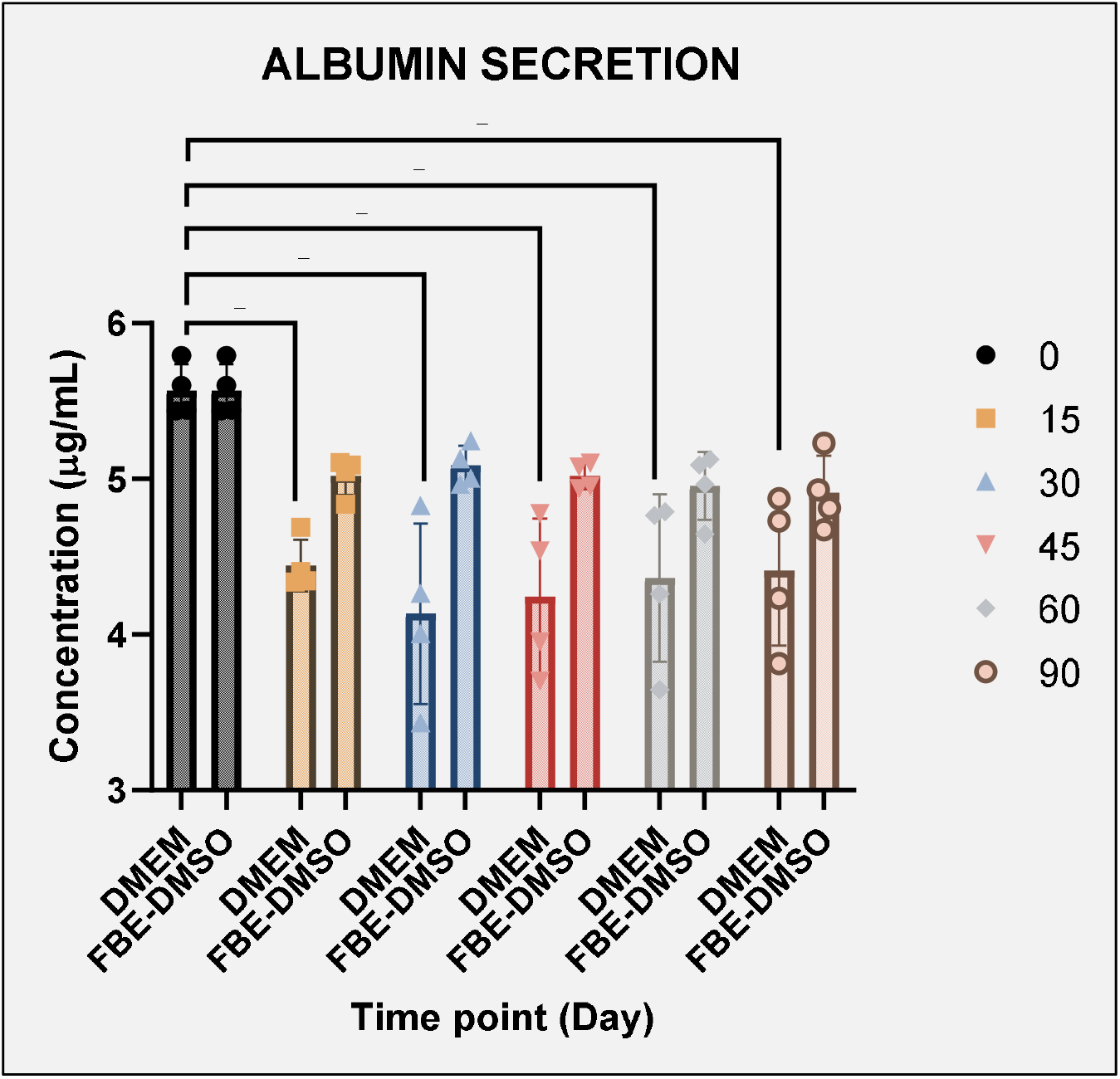
Liver functional assay (A) 3D bioprinted scaffolds stored in DMEM media and (B) 3D bioprinted scaffolds stored in the DMSO and FBS cocktail ; Data presented as mean ± SD (n = 4) statistically significant * : p<0.05 and ns: non-significant

## 4.0 CONCLUSION

This study shows that 3D bioprinted GelMA–dECM liver scaffolds can be stored at −80□°C while still preserving their structure and essential liver functions, provided that a suitable cryoprotectant are employed. When the 3D bioprinted scaffolds were preserved in regular DMEM, there was a steady drop-in the metabolic activity, fewer live cells, and a clear reduction in albumin secretion over 90 days, emphasizing how vulnerable these hydrated 3D bioprinted scaffolds are to damage if the preservation strategy is not optimal. In contrast, 3D bioprinted scaffolds stored in an FBS–DMSO cocktail maintained around 80□% cell viability at all time points and showed stable albumin release, suggesting that this approach effectively protects the cells from ice□ and osmotic□related injury in complex 3D bioprinted liver scaffolds.

The combination of GelMA with rat liver□derived dECM created a more liver□like microenvironment that, together with the optimized cryoprotectant, helped to sustain basic hepatic function post thawing and ready□to□use, cryopreserved 3D liver models. By focusing on post□fabrication preservation, this work begins to close an important gap in tissue□engineering workflows and offers an initial, evidence□based strategy for the long□term storage of 3D bioprinted liver scaffolds. At the same time, the dependence on specific preservation conditions makes clear that further refinement of cryoprotectant formulations, cooling and thawing profiles, and inclusion of more detailed functional and molecular readouts will be essential to develop robust, scalable protocols suitable for industrial translation in drug testing and disease modelling.

## 5. ACKNOWLEDGEMENTS

The authors wish to acknowledge and thank the Department of Biotherapeutics Research (DBR), Manipal Academy of Higher Education (MAHE), and Manipal for the provision of research instruments and infrastructure support.

## 7. AUTHOR CONTRIBUTIONS

Conception and design: Kirthanashri S Vasanthan

Analysis and interpretation of data: Mrunmayi Ashish Gadre, Kirthanashri S Vasanthan Drafting of the paper: Mrunmayi Ashish Gadre and Kirthanashri S Vasanthan

Critical review for important intellectual content: Kirthanashri S Vasanthan

The authors collectively assume responsibility for the integrity and accuracy of all aspects of the research presented.

## 8. DATA AVAILABILITY

Data available within the article or its supplementary materials and the authors confirm that the data supporting the findings of this study are available within the article [and/or] its supplementary materials.

## 10. AVAILABILITY OF DATA AND MATERIALS

The datasets generated and/or analyzed during the current study are available online as the Indian patent, repository (Indian Patent No. 202341045531) and previously published articles with DOIs: https://doi.org/10.1007/s40883-025-00481-2 and https://doi.org/10.1039/D5RA05955K

## 11. ETHICAL APPROVAL

The current research has used animals and was performed with approval of Institutional Animal Ethics Committee (IAEC), protocol number: IAEC/KMC/121/2022, at Manipal Academy of Higher Education, Manipal and under strict adherence to the ARRIVE guidelines, all the protocols including animals were carried out to ensure ethical and humane treatment throughout the research process.

